# Genomic bases underlying the adaptive radiation of core landbirds

**DOI:** 10.1101/2020.07.29.222281

**Authors:** Yonghua Wu, Yi Yan, Yuanqin Zhao, Li Gu, Songbo Wang, David H. Johnson

**Affiliations:** School of Life Sciences, Northeast Normal University, Changchun, 130024, China; Bio-interlligence Co.ltd, Shenzhen 518000, China; Director - Global Owl Project, 6504 Carriage Drive, Alexandria, Virginia 22310 USA

**Keywords:** gene capture, convergent evolution, raptor, digestion, ancestral trait reconstruction

## Abstract

Core landbirds undergo adaptive radiation with different ecological niches, but the genomic bases that underlie their ecological diversification remain unclear. Here we used the genome-wide target enrichment sequencing of the genes related to vision, hearing, language, temperature sensation, beak shape, taste transduction, and carbohydrate, protein and fat digestion and absorption to examine the genomic bases underlying their ecological diversification. Our comparative molecular phyloecological analyses show that different core landbirds present adaptive enhancement in different aspects, and two general patterns emerge. First, all three raptorial birds (Accipitriformes, Strigiformes, and Falconiformes) show a convergent adaptive enhancement for fat digestion and absorption, while non-raptorial birds tend to exhibit a promoted capability for protein and carbohydrate digestion and absorption. Using this as a molecular marker, our results show relatively strong support for the raptorial lifestyle of the common ancestor of core landbirds, consequently suggesting a single origin of raptors, followed by two secondary losses of raptorial lifestyle within core landbirds. In addition to the dietary niche, we find at temporal niche that diurnal birds tend to exhibit an adaptive enhancement in bright-light vision, while nocturnal birds show an increased adaption in dim-light vision, in line with previous findings. Our molecular phyloecological study reveals the genome-wide adaptive differentiations underlying the ecological diversification of core landbirds.

## Introduction

Core landbirds (Telluraves) is the recently defined largest clade in Neoaves, including raptors, woodpeckers, parrots, and songbirds ^1–3^. Core landbirds adapt to diverse environments with different phenotypes related to diets, diel activity patterns, language and hearing, etc. ^4^. Adapting to diverse environments, different core landbirds may have been subjected to divergent selections, whereas for ecologically similar taxa, for instance, three birds of prey (Accipitriformes, Strigiformes, and Falconiformes), which share many raptorial-lifestyle characteristics, similar selections may have occurred between them ^5^. Despite their ecological similarities, phylogenetic analyses show that the three birds of prey are relatively distantly related, and it remains unknown whether their similar raptorial lifestyles are a result of single origin ^2,3^ or multiple origin ^1^.

The Avian Phylogenomics Project evaluates the genomic evolution of 48 avian species within the class Aves with relatively little detailed analyses of genomic difference among avian subgroups, such as core landbirds ^6^. Other comparative genomic or transcriptomic studies related to core landbirds are restricted to specific taxa ^7–13^ or specific genes related to vision and digestion ^14–16^. A recent study of 16 genomes of birds of prey reveals their shared genetic adaptation underlying their anatomical structures and sensory, muscle, circulatory, and respiratory systems related to a raptorial lifestyle ^5^. Despite these efforts, a more comprehensive study with a sufficient sampling across both different taxa and functionally different genes is needed to better elucidate the integrative evolution of core landbirds in adapting to diverse ecological environments.

The recent development of a molecular phyloecological (MPE) approach, which employs a comparative phylogenetic analysis of biologically meaningful genes, has been demonstrated to have power to reveal the genetic bases that underlie phenotypic differentiations and enable ancestral trait reconstruction ^17,18^. In light of the MPE approach, we in this study use target enrichment sequencing to obtain gene sequences to examine the genetic bases that underlie the ecological diversification of core landbirds. Target enrichment sequencing has been widely used in gene capture across species ^19–22^. Our comparative analyses show that different core landbird taxa exhibit taxon-specific adaptations, whereas ecologically similar taxa present somewhat similar evolutionary adaptations. Our study reveals the genomic bases underlying the adaptive radiation of core landbirds and provides insights into the evolutionary history of their ecological diversification.

## Results

We used target enrichment sequencing to obtain the coding sequences of 308 genes related to vision, hearing, language, temperature sensation, beak shape, taste transduction, and carbohydrate, protein and fat digestion and absorption (Table S1, Supplementary text) from 46 core landbird taxa with an address on the sampling of birds of prey (Fig. 1, Table S2), for which there are only sparse genome data available thus far. For the target enrichment sequencing of our focal 308 genes, we designed baits based on the coding sequences of these genes from multiple divergent species (e.g., falcon, eagle and owl), which has been demonstrated to be a highly effective strategy for increasing capture efficiency 22. We then employed the same set of baits to capture and sequence our target genes across the 46 taxa using the MYbaits target enrichment system (MYcroarray, Arbor Biosciences) (Table S2). The average number of reads sequenced across our 46 taxa was 4,716,979.58, the average total base pair yield was 1,178.65 Mb, and the average sequencing coverage and sequencing depth across the 46 taxa were 94.53% and 272.76 X, respectively, while 13 genes showed a sequencing coverage of < 60%, and 17 genes showed a sequencing depth of < 10 X. Raw reads were filtered, and de novo assembled, and coding sequences were obtained for subsequent adaptive evolution analyses. We followed the MPE approach and used likelihood ratio tests based on branch and branch-site models implemented in the Codeml program of PAML 23 to identify positively selected genes (PSGs) along branches leading to diverse core landbird taxa. In addition to PAML, we employed the program RELAX 24 to examine the selection strength changes (e.g., intensified or relaxed) of genes for our focal taxa.

**Fig.1.**
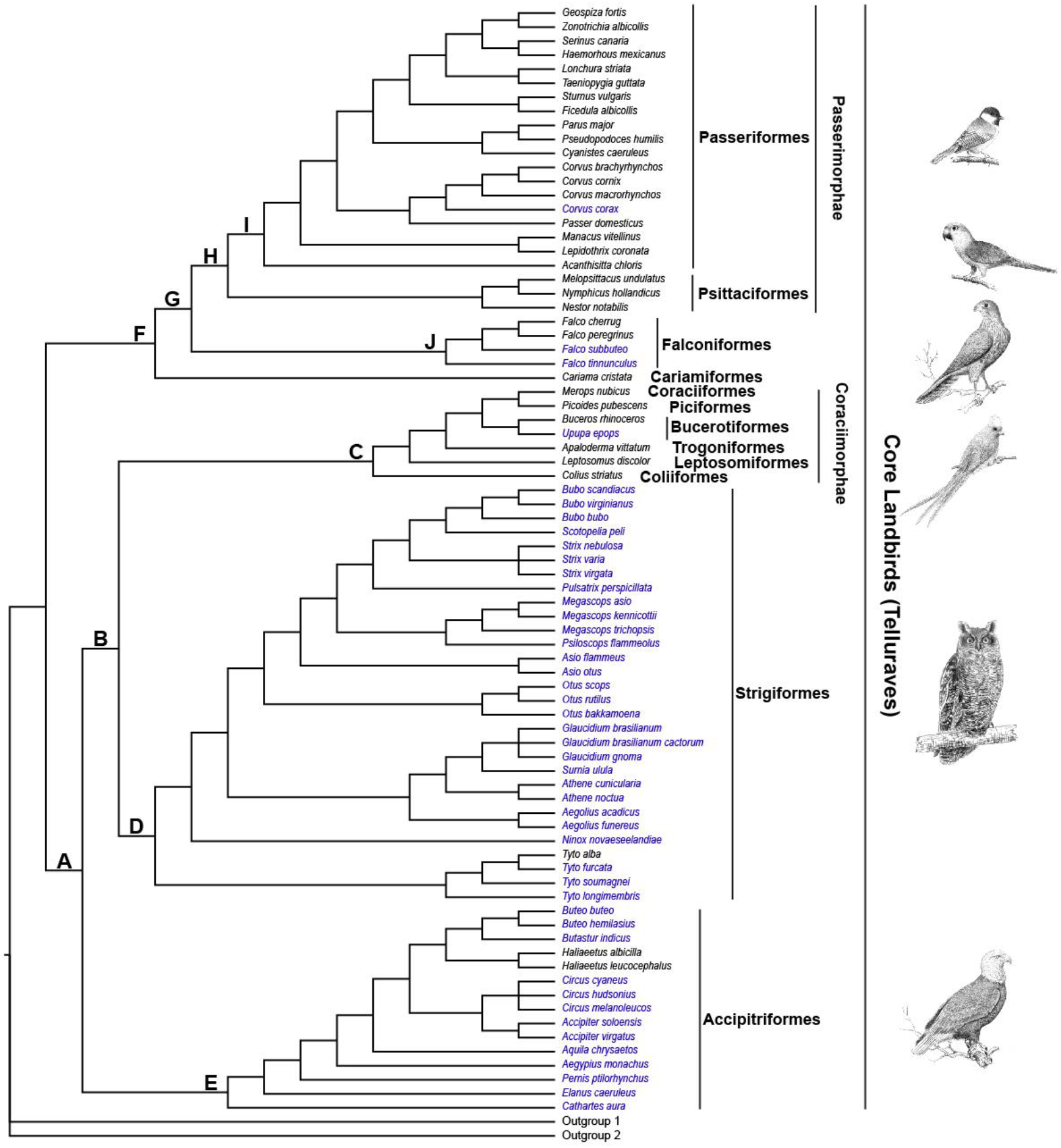
Phylogenetic relationships of core landbird species used in this study. Phylogenetic relationships among species follow published studies^2,17,57,58,62–71^ and the Tree of Life Web Project (http://tolweb.org/tree/). The species under gene capture in this study are shown in blue, and the species with gene sequences cited from GenBank are shown in black. The letters (A-J) show the branches under positive selection analyses based on core landbird dataset in this study.

### Selection analyses of the genes related to senses, language, beak and temperature

To examine the molecular bases of phenotypic differentiations of core landbird taxa, we initially constructed a dataset by combining the captured coding sequences of our 46 taxa with the sequence data of 33 core landbird species available from GenBank (Fig. 1, Table S3), referred to as the core landbird dataset hereafter. Based on the core landbird dataset, we firstly used PAML to examine the adaptive evolution of the 33 genes involved in the phototransduction pathway (Fig. 2, Tables S4-5), which have been used as the molecular markers to determine the diel activity patterns of birds and mammals ^17,18^. We found five PSGs (*GRK1*, *LWS*, *RH2*, *SWS2*, and *ARR3*) along the branch leading to Strigiformes (owls), which are mainly crepuscularly and nocturnally active. *GRK1* is a rod-expressed gene and is involved in the inactivation of activated rhodopsin, contributing to photoresponse recovery in dim-light conditions. *LWS*, *RH2*, and *SWS2*, respectively, encode red-, green-, and blue-sensitive cone opsins, and their positive selections in owls are linked to spectral tuning to maximize light abortion under crepuscular conditions, as previously evidenced 16. *ARR3* is involved in the inactivation of cone opsins for photoresponse recovery. Moreover, we did not find any captured sequences of violet- or ultraviolet-sensitive opsin gene (*SWS1*) from all 28 owl species examined, and this was consistent with previous studies that consistently show the absence of the gene expression or sequence of *SWS1* in owls ^8,9,14,16^. In addition to Strigiformes, we found PSGs along the branch leading to Coraciimorphae and Passeriformes, both of which are generally diurnal ^4,17^. For Coraciimorphae, two PSGs (*CNGB3* and *SLC24A2*) were found. *CNGB3* and *SLC24A2* are cone-expressed and are respectively involved in photoresponse and photoresponse recovery in bright light conditions. For Passeriformes, five PSGs (*GNAT2*, *LWS*, *GUCA1C*, *SLC24A2*, and *RCVRN*) were detected, and almost all of them were cone-expressed, involved in photoresponse and photoresponse recovery in bright light conditions. Finally, we also found two rod-expressed PSGs (*GRK1* and *SLC24A1*) along branch A (Fig. 1), representing the common ancestor of Accipitriformes, Strigiformes and Coraciimorphae.

**Fig.2.**
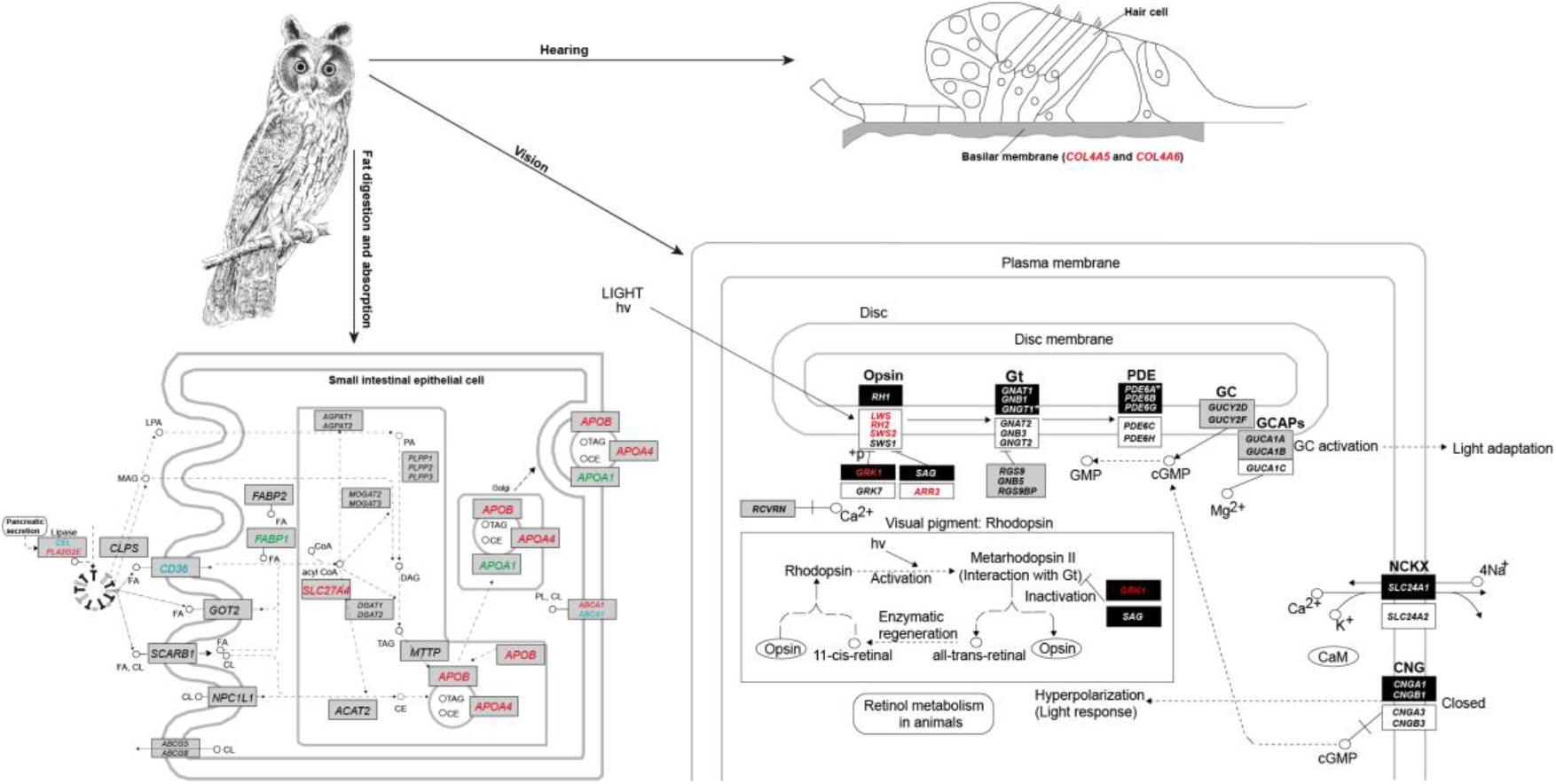
Positively selected and selectively intensified genes related to vision, hearing, and fat digestion and absorption in Strigiformes (highlighted in red). For convenience, the positively selected and/or selectively intensified genes of fat digestion and absorption found in the branches related to Accipitriformes (green) and Falconiformes (blue) are also shown. The top right shows the cross section of inner ear (cochlea), and the lower right shows the phototransduction pathway (according to KEGG pathway map 04744), and the lower left shows the fat digestion and absorption pathway (according to KEGG pathway map 04975). For the phototransduction pathway, the genes that are involved in rods, cones, and both are shown as dark rectangles, white rectangles, and gray rectangles, respectively ^16,18^. *Represents two lost rod-expressed genes, GNGT1 and PDE6A, in both reptiles and birds based on previous studies ^16,18^.

We subsequently examined the positive selection of 95 hearing-related genes, and relevant gene functions were referred to OMIM in NCBI. Our results showed that Coraciimorphae and Passeriformes showed the highest number of PSGs among the diverse core landbird taxa examined (Tables S4-5). For Coraciimorphae, five PSGs (*ATP2B2*, *MYO7A*, *PRPS1*, *SLC9A9*, and *USH2A*) were found. *ATP2B2* plays a critical role in intracellular calcium homeostasis, and its mutation is linked to deafness and imbalance. *MYO7A* plays a role in the development and maintenance of stereocilia, which is critical for converting sound waves to nerve impulses. *PRPS1* encodes an enzyme called phosphoribosyl pyrophosphate synthetase 1, and its mutations have been linked to a type of nonsyndromic X-linked sensorineural deafness. *USH2A* encodes usherin and is an important component of basement membranes in the inner ear and in the retina. *SLC9A9* encodes a sodium/proton exchanger and may play an important role in maintaining cation homeostasis. For Passeriformes, five PSGs (*CRYM*, *FKBP14*, *PTPRQ*, *WNT8A*, and *MYO7A*) were detected. *CRYM* encodes a crystallin protein, and its mutations may cause autosomal dominant non-syndromic deafness. *FKBP14* gene encodes a protein called FKBP prolyl isomerase 14, and its mutations are linked to hearing loss. *PTPRQ* encodes a member of the type III receptor-like protein-tyrosine phosphatase family, and its mutations may cause autosomal recessive deafness. *WNT8A* encodes a protein of the WNT gene family and may be involved in the development of early embryos and germ cell tumors. In addition to Coraciimorphae and Passeriformes, positively selected hearing-related genes were found in other taxa. Notably, *COL4A5* and *COL4A6* were found to be under positive selection in owls (Fig. 2, Table S5). The two genes encode two of the six subunits of type IV collagen, the major structural component of basement membranes in the inner ear. Mutations of the two genes are associated with hearing loss and the severity of deafness. For Falconiformes (falcons), two PSGs (*MYO7A* and *SLC17A8*) were detected; *MYO7A* is involved in the development and maintenance of stereocilia, and *SLC17A8* encodes a vesicular glutamate transporter. We also detected one positively selected gene, *TMC1*, along branch H, which represents the common ancestor of Passeriformes and Psittaciformes. *TMC1* is required for the normal functioning of cochlear hair cells, and its mutations have been associated with progressive postlingual hearing loss and profound prelingual deafness.

We also analyzed the positive selection of the genes related to taste, language, beak shape, and temperature sensation (Tables S4-5). Among the 43 taste-related genes analyzed, several PSGs were found in different taxa. Though these PSGs found are related to taste transduction, only one positively selected gene, *SCNN1A*, which was found along the branch leading to Accipitriformes, is relatively clearly related to taste (salt) perception; other PSGs, such as *PDE1C*, *GNB3*, and *ENTPD2*, are found to be widely expressed in many other tissues as well (please see OMIM in NCBI for details). We also examined the positive selection of 24 language- and/or speech-related genes, and our results showed that *ROBO1* in owls, *DCDC2* in falcons, and *FOXP2* in the common ancestor of Falconiformes and Passerimorphae were found to be under positive selection. *ROBO1* is involved in core trait underpinning language acquisition, *DCDC2* encodes a doublecortin domain-containing family member and is linked to dyslexia, and *FOXP2* encodes a member of the forkhead/winged-helix (FOX) family of transcription factors and is required for the proper development of speech and language. Finally, we analyzed the positive selection of 15 beak shape-related genes and 15 temperature sensation-related genes. For the beak shape-related genes, we found only one positively selected gene, *DKK3*, which showed a positive selection signal in owls. *DKK3* plays a role in the development related to beak depth and length. For the temperature sensation-related genes, three PSGs were detected, including one cold sensor (*TRPA1*) in owls, one warm sensor (*TRPM2*) in falcons, and one warm sensor (*TRPV4*) in the common ancestor of owls and their sister taxon, Coraciimorphae.

### Selection analyses of digestive system-related genes

We used PAML to examine the adaptive evolution of the 84 genes that are involved in the carbohydrate, protein and fat digestion and absorption pathways to determine the possible selection differences between core landbird taxa. Our initial analyses using PAML showed that almost all examined branches exhibited positive selection signals in the genes related to carbohydrate and/or protein digestion and absorption to some extent, with the exception of two raptor-related branches (branches G and D, Fig. 1), which also showed positive selection on the genes related to fat digestion and absorption, including one gene (*CEL*) along the common ancestor branch (branch G) of Falconiformes and Passerimorphae, and three genes (*APOA4*, *APOB*, and *SLC27A4*) along the branch (branch D) leading to owls (Tables S4-6). To further examine whether the positive selections of the fat digestion and absorption-related genes occurred uniquely along raptor-related branches, we then respectively evaluated the relative selection intensity changes of the 84 genes along the branches leading to three birds of prey and their sister taxa compared to their common ancestor branches using program RELAX ^24^ (Fig. 1, Table S7). Intriguingly, our results showed that all three birds of prey showed selectively intensified genes related to fat digestion and absorption (Fig. 2), including *CD36* in Falconiformes (branch J compared to branch G), *ABCA1* in Strigiformes (branch D compared to branch B), and *APOA1* in Accipitriformes (branch E compared to branch A), though Falconiformes and Strigiformes showed a selection intensification on the genes related to carbohydrate and/or protein digestion and absorption as well (Tables S6-7). Unlike the three birds of prey, which almost consistently exhibited the selection intensification of the genes related to fat digestion and absorption to some extent, their sister taxa, Coraciimorphae and Passerimorphae, showed somewhat variable results (Tables S6-7). Especially for Coraciimorphae (branch C), almost all detected genes showed a relative selection relaxation compared to the common ancestor branch (branch B) of Coraciimorphae and Strigiformes, and these genes were found to be mainly related to fat and protein digestion and absorption. For Passerimorphae (branch H), four genes (*CD36*, *CPB2*, *CTRL* and *PRKCB*), which are involved in fat, protein and carbohydrate digestion and absorption, showed a relative intensified selection compared to the common ancestor branch (branch G) of Passerimorphae and Falconiformes, while two genes (*LCT* and *PIK3CD*) related to carbohydrate digestion and absorption showed a relative selection relaxation. Taken together, it seems that all three birds of prey showed a relative intensified selection for fat digestion and absorption to some extent, while their non-raptorial relatives showed somewhat variable results.

Our RELAX results described above revealed the selection differences of the digestive system-related genes between raptors and non-raptorial birds, showing that all three birds of prey were characterized by an intensified selection for fat digestion and absorption. In light of this finding, we subsequently examined the selection intensity changes of these digestive system-related genes for the common ancestor of core landbirds studied, which is considered to be an apex predator ^2,3^. To this end, we constructed a new dataset, which included largely the full-length coding sequences of these digestive system-related genes from all avian taxa with gene sequences available from GenBank (Fig. 3, Table S8), referred to as the whole avian dataset hereafter, considering that our core landbird dataset used involved only a few outgroups of core landbirds (Fig.1), and that its gene sequences used were relatively short. Based on the whole avian dataset, we examined the relative selection intensity changes of our focal genes along the common ancestor branch (branch L) of core landbirds compared to the common ancestor branch (branch M) of core landbirds and their sister taxa, waterbirds (Fig. 3). Our results showed that the common ancestor of core landbirds exhibited a marked intensified selection on the two genes (*APOA4* and *FABP1*) related to fat digestion and absorption (Table S6, Table S9), with a much larger K value (K ≈ 50, p < 0.05) than a neutrality indicator of K = 1. *APOA4* encodes apolipoprotein A4 and is a potent activator of lecithin-cholesterol acyltransferase. *FABP1* encodes the fatty acid binding protein found in liver and plays a role in fatty acid uptake, transport, and metabolism (Table S6). We also found that another two fat digestion and absorption-related genes (*AGPAT1* and *CD36*) showed a relatively slight selection relaxation, with K values of 0.95 and 0.89, respectively. Moreover, we found three carbohydrate digestion and absorption-related genes (*HK3*, *PIK3CD*, and *SI*) were under an intensified selection, and two protein digestion and absorption-related genes (*MME* and *SLC9A3*) respectively showed a slightly intensified selection and a marked selection relaxation (Table S9). It seems that the ancestral core landbirds were subject to a relatively intensified selection for fat and carbohydrate digestion and absorption, which was more similar to that found in owls (Tables S4-7).

**Fig.3.**
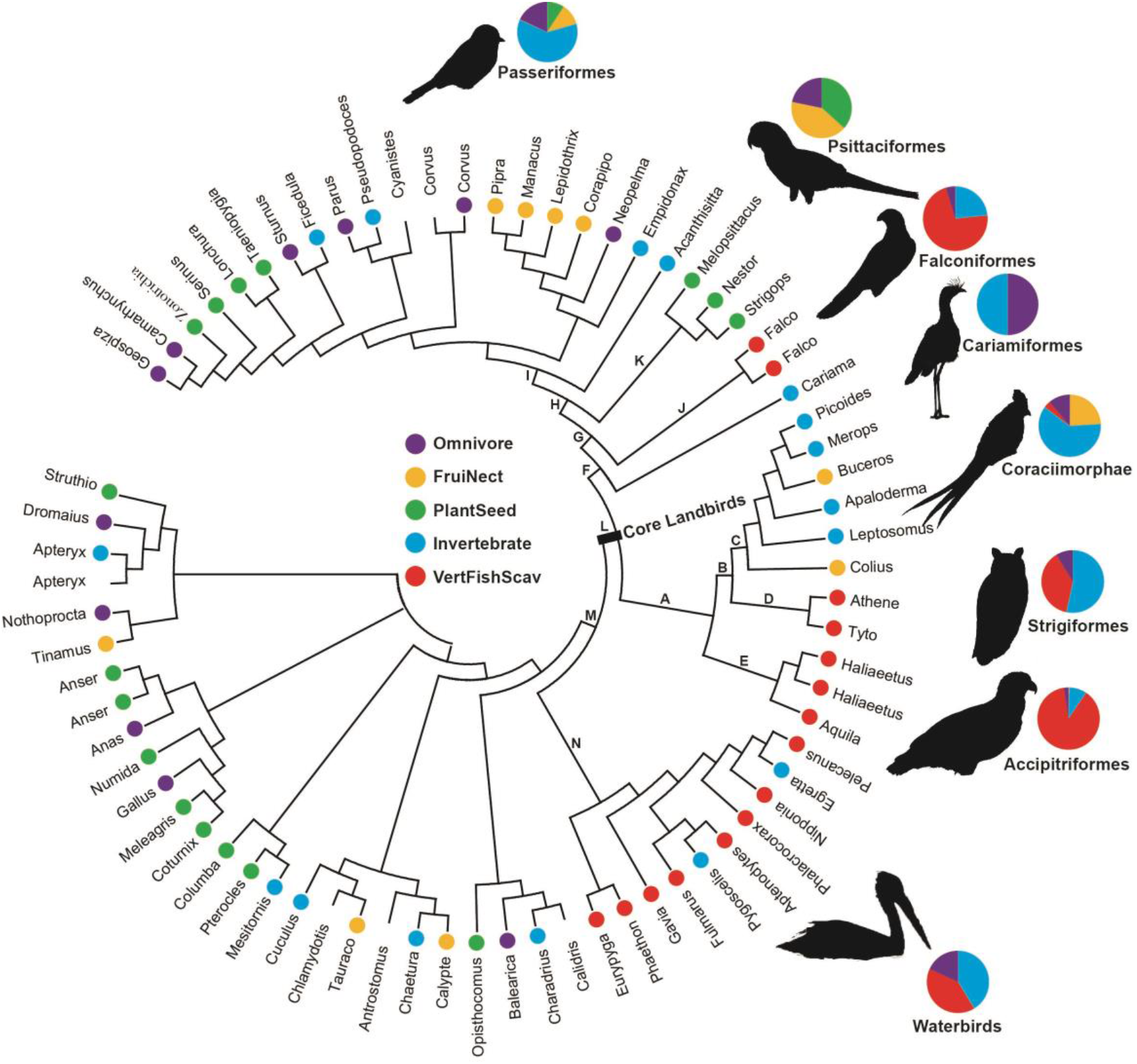
Avian phylogeny and dietary categories. Avian phylogeny is constructed based on published studies 17,67,69 and the Tree of Life Web Project (http://tolweb.org/tree/). For convenience, only the genus names of species are shown. Dietary information is derived from a previous study ^59^. Dietary categories are shown in different colors, and blanks indicate that dietary information is unavailable. The pet charts show the proportions of species numbers with different dietary categories for our focal avian groups. PlantSeed (plant and seeds), FruiNect (fruits and nectar), Invertebrate (invertebrates), VertFishScav (vertebrates and fish and carrion), and Omnivore (score of <= 50 in all four categories). The letters (A-N) show the branches under positive selection analyses based on the whole avian dataset in this study.

Based on the whole avian dataset, we also conducted the positive selection analyses along the common ancestor branch of core landbirds, as well as along other branches of interest (branches A-N) (Fig. 3, Tables S10-11). No PSGs were found along the common ancestor branch of core landbirds, while we found that all three birds of prey harbored PSGs that are involved in fat digestion and absorption, and non-raptorial birds showed predominant positive selection on the genes involved in protein or carbohydrate digestion and absorption (Table S6, Tables S10-11). Specifically, for the three birds of prey, two PSGs from each of them were found, with *MEP1B* and *PLA2G2E* found in Strigiformes, *CPB1* and *FABP1* found in Accipitriformes, and *G6PC2* and *ABCA1* found in Falconiformes. Among the six PSGs detected, *PLA2G2E*, *FABP1*, and *ABCA1* are involved in fat digestion and absorption, while the other three are related to either protein or carbohydrate digestion and absorption (Table S11). In addition to the raptors, we also found one PSG (*SLC15A1*) along the Coraciimorphae branch, and one PSG (*G6PC3*) along the Passerimorphae branch, and six PSGs (*CELA3B*, *SLC15A1*, *CTRL*, *HK3*, *G6PC3*, and *APOB*) along Passeriformes branch. However, almost all of these PSGs found in the non-raptorial birds are involved in protein or carbohydrate digestion and absorption, with the exception of *APOB* (Table S11). We also examined the positive selection along the branch leading to parrots (Psittaciformes), and strikingly, we found 8 PSGs. Among the eight PSGs, four (*APOA4*, *APOB*, *CLPS*, and *ABCG5*) are involved in fat digestion and absorption, three (*MME*, *CPA2*, and *PRCP*) are involved in protein digestion and absorption, and one (*HK3*) is involved in carbohydrate digestion and absorption. Finally, we examined the positive selection along the common ancestor branch (branch M) of core landbirds and their sister taxa, waterbirds (Fig. 3, Tables S10-11), and one gene, *AGPAT1*, which is involved in fat digestion and absorption (Table S6), was found to be under positive selection. Taken together, our results based on both the core landbird dataset and the whole avian dataset consistently show that the three birds of prey exhibited an enhanced selection for fat digestion and absorption, while their non-raptorial relatives tend to exhibit an enhanced selection for protein and/or carbohydrate digestion and absorption.

## Discussion

We in this study used target enrichment sequencing to obtain the coding sequences of 308 genes to examine the molecular bases that underlie the ecological diversification of core landbirds in the context of the MPE approach. Our results showed that different core landbird taxa exhibit evolutionary enhancements in different aspects, suggesting lineage-specific selection. For instance, for owls, which are mainly crepuscularly and nocturnally active and are well known to have acute hearing ^25^, we detected evidence of the positive selection of five vision genes that mainly contribute to dim-light vision (Fig. 2, Table S5) and their absence of the *SWS1* sequence, suggesting enhanced dim-light vision and decreased bright-light vision, consistent with previous studies ^5,9,16^. Besides their enhanced dim-light vision, we also found evidence of the positive selection of two hearing-related genes, *COL4A5* and *COL4A6*, in owls (Table S5). The two genes play a critical role in formation of basement membranes (Fig. 2), which are responsible for transferring sound waves to the organ of Corti for the sense of hearing. The positive selection of the two hearing-related genes in owls may suggest their increased sensitivity to sound in dim-light conditions. Moreover, we also detected evidence of the positive selection of one cold-sensor gene, *TRPA1*, in owls (Table S5), which may be useful for them to sense low ambient temperatures at night. Unlike owls, for two diurnal taxa, Coraciimorphae and Passeriformes, we found almost solely cone-expressed vision genes, which are involved in photoresponse and photoresponse recovery, to be under positive selection (Tables S4-5), suggesting their increased visual acuity and promoted capacity for motion detection in bright-light conditions. In addition to their promoted bright-light vision, we also detected their predominant positive selections on hearing-related genes, suggesting that they may have evolved enhanced hearing as well. Many members of Coraciimorphae and Passeriformes are arboreal and are very vocal ^4^, and a promoted bright-light vision and enhanced hearing may help in the avoidance of obstacles and facilitating vocal communication in forest environments. We also detected the positive selection of language-related genes in owls and falcons, and particularly, we found the positive selection of the *FOXP2* gene, a well-known language gene, in the common ancestor of falcons and Passerimorphae (Table S5), suggesting their evolutionary modification of language.

Our results demonstrated that all three birds of prey convergently evolved an enhanced capability for fat digestion and absorption, while non-raptorial birds tend to exhibit a promoted evolutionary enhancement for protein and carbohydrate digestion and absorption (Tables S6-7, Tables S10-11). Consistent with our finding of an increased capability for fat utilization in birds of prey, one recent study shows the relatively accelerated evolution of one lipid metabolism-related gene, *PPARA*, across birds of prey ^5^. Previous studies show that the digestive physiology of animals evolves in parallel with their diets ^26^, and their digestion enzyme activities are known to be positively correlated with the amount of protein, carbohydrate, and lipid intake ^27^. Our finding of an enhanced capability for fat utilization in birds of prey may be associated with their specialized carnivorous diets. All three birds of prey consume a considerable amount of vertebrates as prey (Fig. 3), which are relatively rich in protein and fat ^28^. Fats contain the highest amounts of energy, with more than twice the number of calories as the same amount of proteins or carbohydrates^28^. An enhanced capability for fat utilization may help birds of prey to adapt to their relatively fat-rich diet. Unlike birds of prey, non-raptorial birds, such as Passeriformes and Coraciimorphae, eat a considerable amount of invertebrate as prey as well as plant food (e.g., seeds, fruits and nectar) (Fig. 3), which are generally protein-rich and carbohydrate-rich, respectively ^28–30^. An increased capability for the utilization of protein and carbohydrates in non-raptorial birds may be partly attributed to the relative high proportion of protein and carbohydrates in their diets. Consistent with this, it has been known that most herbivores, from fishes to mammals, have a very high number of glucose transporters relative to carnivores, suggesting a relative evolutionary enhancement of carbohydrate utilization in herbivores ^26^.

Consistent with the enhanced capability of lipid utilization in birds of prey found in this study, previous genomic studies have well documented an enhanced lipid utilization in many carnivorous mammals (e.g., dog, cats, polar bear, cetaceans) ^31–40^, carnivorous reptiles (e.g., pythons, Komodo dragons) ^41,42^ and even modern humans, such as the Maasai, who consume a fat-rich diet ^43^. In addition to these findings at the molecular level, anatomically, carnivores are usually found to retain a gallbladder, and their gallbladders are relatively large, while for those animals that feed primarily on plant food, the gallbladder is absent or relatively small ^26,44^. The gallbladder is used for storing bile, which is essential for the digestion of fat. The evolutionary enhancement of gallbladders in carnivorous animals may suggest a relatively high selection pressure for fat digestion, in line with the molecular findings mentioned above. Nevertheless, an enhanced lipid metabolism has also been reported in some ruminants (e.g., sheep, deer), which are typically herbivores. However, this is considered to be mainly due to their dependence on volatile fatty acids (VFAs) as an important nutrient source, which are generated by their rumen microbial flora from plant material ^45,46^. It should be noted that we also detected a notable evolutionary enhancement of fat utilization-related genes along the branches leading to Passerimorphae (parrots to songbirds) and parrots, respectively (Table S7, Table S11). We speculated that this may be partly attributed to the specific diet of parrots. Parrots largely include plant food, including nuts and seeds ^47^, in their diets (Fig. 3), and in particular, they have evolved a short, deep, hooked beak and a shortened, downward-curved lower jaw, which are adapted for cracking hard nuts and seeds 48, and nuts are generally high in fat ^28^. One recent study shows that 37.68% of parrot species are known to ingest nuts in their diet, and these species are scattered across almost the whole parrot phylogeny, including the deepest and basal branches ^47^ (Fig. S1). This may suggest that an increased capability of fat utilization may have evolved early accompanying the origin of parrots. These findings may suggest that the digestive system-related genes of animals may have been evolutionarily modified to adapt to the nutrient compositions (e.g., proteins, carbohydrates, and lipids) that they directly utilize in their diets.

The differentiated selections of digestive system-related genes between raptors and non-raptorial birds found in this study shed light on the origin of raptorial lifestyles in core landbirds. Core landbirds harbor three relatively distantly related birds of prey (Accipitriformes, Strigiformes and Falconiformes), and previous studies hypothesize that the common ancestor of core landbirds may be an apex predator ^2,3^. Our findings show that the common ancestor of core landbirds exhibited a particularly enhanced selection intensification for fat digestion and absorption (Table S9). And this is similar with that observed in the three birds of prey, providing a relatively strong support for this hypothesis. However, we could not completely exclude the possibilities that the common ancestor of core landbirds could be of other types of dietary guilds, such as a plant eater that relies on a fat-rich plant food (e.g., nuts). However, other lines of evidence provide additional support for the common ancestor of core landbirds as an apex predator, as follows: 1) The three birds of prey are phylogenetically positioned at the deepest and/or basal branches within core landbirds ^2,3^, suggesting their possibly early evolution. 2) Fossil evidence indicates that birds of prey (owls) occur earlier than almost all other members of core landbirds ^49,50^. 3) Our selection analyses demonstrate that Coraciimorphae showed a substantial selection relaxation of fat and protein digestion and absorption-related genes (Table S7). This is consistent with their possible secondary loss of the raptorial lifestyle. 4) Previous studies demonstrate high transition rates from a carnivorous diet to other dietary guilds but less occurrences vice versa during the evolutionary histories of birds and mammals ^51,52^. This is consistent with the proposed raptorial lifestyle of ancestral core landbirds and their subsequent evolutionary transitions to the two non-raptor lineages leading to Passerimorphae and Coraciimorphae ^2^.

Core landbirds may have initially evolved in South America ^53^ as an apex predator, closely related to the extinct top predators, Phorusrhacidae, in the Cenozoic of South America ^54,55^, and subsequently they dispersed globally and underwent an adaptive radiation to make use of different niches ^53^. Accipitriformes hunt by day, Strigiformes hunt by night, while Falconiformes hunt by both day and night ^16^. And two lineages, Passerimorphae and Coraciimorphae, may be secondarily derived from their raptorial ancestors and are mainly active during the day, as evidenced in this study, and evolved to exploit other non-raptorial diet niches (Fig. 3) ^2,54^. Accumulated fossil evidence has shown that Eocene relatives of Coraciimorphae members (mousebirds and cuckoo-rollers) and Passerimorphae (parrots) have raptor-like feet or claws, and stem group passerines show parrot-like skeletal morphology ^2,56^. These transitional fossils may suggest a gradual evolutionary change of Coraciimorphae and Passerimorphae from their raptorial-like ancestors ^2^. If this is the case, we may speculate that the resemblance of parrots to birds of prey, such as their raptor-like curved beak, may be derived from their raptorial progenitors as well. It is noteworthy that our finding of the positive selection of one fat utilization-related gene, *AGPAT1*, along the common ancestor branch of core landbirds and their sister taxa, waterbirds, suggests the possibility of its raptorial lifestyle. And this is consistent with the considerable occurrences of carnivorous birds that prey on vertebrates in both core landbirds (e.g., birds of prey) and waterbirds (Fig. 3). This may imply that a raptorial lifestyle may have evolved much earlier than previously realized.

## Conclusion

Our study shows that core landbirds are subject to lineage-specific selections, and that different groups show evolutionary enhancements in different aspects. Despite this evolutionary specialization, two general evolutionary patterns emerge. First, nocturnal birds are characterized by an evolutionary enhancement in dim-light vision, while diurnal birds tend to evolve an enhanced bright-light vision, in line with previous studies ^16^. More importantly, we find that the adaptive evolutions of digestive system-related genes are capable of reflecting the changes of nutrient compositions (e.g., proteins, carbohydrates, and lipids) in the diets of different avian taxa. This provides an important insight into reconstructing the ancestral dietary state of animals in the context of molecular phyloecology ^17,18^. Particularly, our results based on two different datasets (core landbird dataset and whole avian dataset) consistently demonstrated that all three birds of prey show a convergent evolutionary enhancement for fat digestion and absorption, while non-raptorial birds tend to evolve an increased capability for protein and carbohydrates digestion and absorption, in relation to their differentiated diets. In light of this finding, our results provide relatively strong support for the raptorial lifestyle of ancestral core landbirds, implying a single origin of birds of prey, followed by two subsequently secondary losses in other non-raptorial birds. Moreover, our study implies that a raptorial lifestyle likely occurred much earlier than previously realized, possibly at least tracing back to the common ancestor of core landbirds and their sister taxa, waterbirds.

## Materials and method

### Datasets and taxa used

In this study, we constructed two datasets, the core landbird dataset (Fig. 1) and the whole avian dataset (Fig. 3). In the core landbird dataset, the coding sequences of 78 core landbird species and a few outgroups were involved. Among the 78 species, 45 species were under target enrichment sequencing in this study, and the other 33 species’ gene sequences were cited from GenBank. The 45 species under the target enrichment sequencing included 28 Strigiformes species, 13 Accipitriformes species, two Falconiformes species, one Coraciimorphae species and one Passeriformes species (Fig. 1). Particularly, the 28 species of Strigiformes were selected to represent all of its two known extant families (Tytonidae and Strigidae), five subfamilies, and nine tribes, with the only exception being Phodilinae ^57^. The 13 species of Accipitriformes were from nine genera, two families (Cathartidae and Accipitridae), and particularly, 12 species of them were from Accipitridae and represented many of its divergent clades ^58^. In addition to the core landbird dataset, which contained species from almost solely core landbirds, we also constructed the whole avian dataset, which contained the full-length coding gene sequences from all avian species (73 species in total) available from GenBank (Fig. 3). In total, 125 avian taxa across the core landbird dataset and whole avian dataset were used in this study.

### Diet dataset

Avian diet information was used from one dataset, EltonTraits 1.0, which includes the compiled dietary information of a total of 9993 extant bird species from diverse published literature ^59^. Accordingly, each species is assigned to the dominant among five diet categories: PlantSeed (plant and seeds), FruiNect (fruits and nectar), Invertebrate (invertebrates), VertFishScav (vertebrates and fish and carrion), and Omnivore (score of <= 50 in all four categories). In addition to EltonTraits 1.0, we also referred to another parrot-specific dataset for its dietary analyses in the context of parrot phylogeny. This dataset includes a time-calibrated phylogenetic supertree of all 398 extant parrot species and detailed information about their diet components (e.g., seeds, nuts) ^47^.

### Gene list used

The genes related to vision, hearing, language, temperature sensation, beak shape, taste transduction, and carbohydrate, protein and fat digestion and absorption were involved in this study (Table S1). In total, 308 genes were included. These gene lists involved in the different biological functions were compiled from the published literature and/or online KEGG Pathway database (please see supplementary text for details). Briefly, for vision, the 33 phototransduction genes that are used to determine diel activity patterns ^17,18^ were used. For hearing, the 95 genes related to hearing were retrieved from the published literature. For temperature sensation, the 15 genes that are mainly involved in thermosensitive TRP channels spanning from sensing warm to cold temperatures were used based on the published literature. For language and beak shape, the related genes were mainly retrieved from the published literature. Finally, for taste transduction, and carbohydrate, protein and fat digestion and absorption, the gene lists were retrieved from the relevant KEGG pathways (maps 04742, 04973, 04974, 04975). The function descriptions of these genes were referred to the published literature and the KEGG Pathway database (Supplementary Text) as well as the Online Mendelian Inheritance in Man (OMIM) and/or Gene in NCBI.

### Bait design

We designed baits mainly based on the coding sequences of 308 genes of three species from GenBank, including *Falco cherrug*, *Haliaeetus albicilla*, and *Tyto alba*, representing three birds of prey, Falconiformes, Accipitriformes, and Strigiformes, respectively. If their gene sequences were unavailable for a certain gene, the gene sequences from their phylogenetical relatives were used, including *Falco peregrinus*, *Aquila chrysaetos*, *Haliaeetus leucocephalus*, *Picoides pubescens*, *Apaloderma vittatum*, and *Colius striatus*. Moreover, we also included the transcript sequences of 33 phototransduction genes from 15 core landbird species published previously in our lab ^16^. Based on these gene sequences, we designed 90nt probes with 2x tiling density, then we screened them against *Falcon peregrinus* genome with high-coverage ^6^ (GCF_000337955.1_F_peregrinus_v1.0_genomic.fna.gz) as well as the owl mitochondrial genome, and we applied an “ultrastringent” level of filtration to identify and remove those baits that were likely to hybridize and capture more than one genomic region, and eventually, 59,969 ultrastringent-baits were used for subsequent target enrichment via hybridization-based capture (MYcroarray, Arbor Biosciences, USA).

### Sample collection, DNA extraction and target enrichment sequencing

We collected blood, muscle, and liver samples from 46 core landbird taxa (Table S2), of which 27 blood samples were obtained in this study, 15 muscle samples were collected previously in our lab 16, and four liver samples were contributed by the Jilin Provincial Museum of Natural History. The 27 blood samples obtained in this study were collected onto FTA cards and then mailed to a laboratory and stored at −80 ℃ condition. We abstracted genomic DNA from the blood samples using the DNeasy Blood & Tissue Kit (QIAGEN), and we abstracted genomic DNA from the muscle or liver samples using the Ezup Column Animal Genomic DNA Purification Kit (Sangon Biotech, Beijing). DNA concentration was measured by NanoDrop 2000C (Thermo Scienfitic) (Table S2).The experimental procedures were performed in accordance with an animal ethics approval granted by Northeast Normal University. All experimental procedures carried out in this study were approved by the National Animal Research Authority of Northeast Normal University, China (approval number: NENU-20080416) and the Forestry Bureau of Jilin Province of China (approval number: [2006]178).

We quantified the DNA samples of the 46 taxa using the Quant-iT(TM) Picogreen(R) dsDNA Assay kit (ThermoFisher). The visualization of a subset of samples on a Bioanalyzer High-Sensitivity DNA Chip (Agilent) indicated that the DNA was predominantly high in molecular weight. Up to 4ug of each sample was then sheared with a Qsonica Q800R instrument and selected to modal lengths of roughly 300nt using a dual-step SPRI bead cleanup. We then converted on average 384 ng of the sheared, size-selected samples to Illumina(R) sequencing libraries using chemistry based on the KAPA HyperPrep(TM) (Kapa Biosystems) kit for Illumina. After ligation, the libraries were amplified using KAPA HiFi HotStart ReadyMix (Kapa Biosystems) for 6, 8 or 10 cycles, inversely scaled to the mass taken to the library preparation, using the manufacturer’s recommended thermal profile. Custom i7 and i5 8nt indices were incorporated during this amplification in combinations designed to prevent cross-indexing during post-capture pooled amplification. After purification with SPRI beads, we then quantified each index-amplified library using the Quant-iT(TM) Picogreen(R) dsDNA Assay kit (ThermoFisher). For target enrichment, two samples (*Corvus corax* and *Upupa epops*) were captured individually using 500ng total library per reaction. All other samples were captured in pools of eight samples, using 80% of the available library mass up to 100 ng each. Enrichment followed the myBaits Manual version 3 using double the block #3 reagent and 65℃ temperatures for all relevant steps. After enrichment, half of the cleaned bead suspension was taken to 10 cycle amplification using universal primers and KAPA HiFi HotStart ReadyMix (Kapa Biosystems). After purification, each captured library was quantified with quantitative PCR and combined in equimolar ratios prior to sequencing. The equilibrated sequencing pool was then submitted to the Genomic Services Lab at Hudson Alpha (Huntsville, Alabama, USA) for paired-end 125 bp sequencing on a HiSeq 2500 instrument using version 4 chemistry. Raw FASTQs were delivered following the standard CASAVA post-processing pipeline, and were deposited in the Sequence Read Archive (SRA) database with accession numbers (SRR11449714-59).

### Sequence assembly and alignment

Raw reads were processed by SOAPnuke (version 1.5.6, https://github.com/BGI-flexlab/SOAPnuke) to remove adapter sequences and low quality reads with the following parameters: -n 0.01 -l 20 -q 0.4 -Q 2. Then, duplicate reads were filtered using a custom Perl script. Prior to assembly, all clean reads were aligned to our target sequences using BLASTN to identified intron-exon boundaries (e value cutoff = 1e^−5^), and then intron sequences were trimmed. All processed reads were de novo assembled into contigs by Trinity ^60^ with the following parameters: --min_kmer_cov 3 --min_glue 3 --bfly_opts ‘-V 5 --edge-thr=0.1 --stderr’ --min_contig_length 150. Then all contigs were aligned to our target gene sequences originally used for bait design using the BLASTP with default parameters. And we chose hits with an identity >80% and coverage > 60% as the candidate contigs that were associated with certain targets (in-target assemblies). Subsequently, these candidate contigs were joined together with Ns based on their relative BLAST hit positions in regard to the target sequences. To verify the orthology of these candidate contigs, they were then blasted against the National Center for Biotechnology Information (NCBI) nr/nt database using BLASTN online. We kept for subsequent analyses only those candidate contigs with their best hit being our target gene; otherwise, that particular sequence was discarded. Finally, sequencing coverage and sequencing depth were evaluated by BWA ^61^ with a default parameter to evaluate capture efficiency. Gene sequences were aligned using webPRANK (http://www.ebi.ac.uk/goldman-srv/webprank/), and individual sequences with low identities, long indels, multiple ambiguous bases Ns, and/or too short a length were removed. After this pruning, high-quality alignments were constructed, and their translated protein sequences were blasted against the non-redundant protein sequence database to confirm the correctness of sequence cutting.

### Positive selection analyses

We used the branch and branch-site models implemented in the Codeml program of PAML ^23^ for our positive selection analyses. For this, we constructed unrooted species trees following previous studies ^2,17,57,58,62–71^ and the Tree of Life Web Project (http://tolweb.org/tree/). We used a codon-based maximum-likelihood method to evaluate the ratio of non-synonymous to synonymous substitutions per site (dN/dS or ω), and used likelihood ratio tests (LRT) to determine statistical significance. A statistically significant value of ω > 1 suggests a positive selection. Specifically, for the branch model analyses, we used a two-rate branch model to detect possible positive selection signals along a particular branch. With the analyses, our focal branches were respectively labeled foreground branches, and others were used as background branches. The two-rate branch model was then compared with the one-rate branch model to determine statistical significance using LRT. If a statistically significant value of ω > 1 in a foreground branch was found, the two-ratio branch model was then compared with the two-ratio branch model with a constraint of ω = 1 in the foreground branch to further examine whether the ω > 1 of the foreground branch was supported with statistical significance. In addition to the branch model, we also used a branch-site model (Test 2) to detect positively selected sites along our focal branches (foreground branches). Test 2 compares a modified model A with its corresponding null model with ω = 1 constrained to determine the statistical significance using LRT. The empirical Bayes method was used to identify positively selected sites. For our result robustness, we further examined the dependency of our positive selection results on the initial value variations of kappa and omega. For this, we respectively used two different initial values of kappa (kappa = 0.5, 3.0) and of omega (ω = 0.5, 2.0) for our positive selection analyses, and in total, four independent runs were conducted for each of the PSGs found.

### Selection intensity analyses

We evaluated the relative selection intensity changes of genes using RELAX ^24^, which is available from the Datamonkey webserver (http://test.datamonkey.org/relax). The relative selection intensity of genes was measured using a parameter k value. The k value and its statistical significance were evaluated given a priori partitioning of test branches and reference branches in a codon-based phylogenetic framework. K > 1 shows an intensified selection, and k < 1 shows a relaxed selection. Intensified selection is expected to exhibit ω categories away from neutrality (ω = 1), while relaxed selection is expected to exhibit ω categories converging to neutrality (ω = 1). Statistical significance was evaluated by comparing an alternative model to a null model using LRT. The alternative model assumes different ω distributions of the test and reference branches, while the null model assumes k = 1 and the same ω distribution of both test and reference branches.

## Acknowledgement

We acknowledge Michael Wink, Russell Thorstrom, Yu Liu, Wei Liang and Haitao Wang for helping sample collections. This research was supported by the National Natural Science Foundation of China (grant number, 31770401) and the Fundamental Research Funds for the Central Universities.

## Author contribution

Y. W. conceived the study, performed experiments, conducted data analyses, and wrote the paper. D.H.J conceived the study, collected blood samples and read the manuscript. S.W helped for sequence assembly and coding sequence abstraction. Y.Y., Y.Z. and L.G. helped for positive selection analyses.

## Supplementary text

### Supplementary figure and table legends

**Fig. S1** Phylogenetic supertree of 413 parrot species. Parrots that consume nuts in their diets are shown in red. The phylogenetic supertree and dietary information follow one published study (Burgio et al. 2019).

**Table S1** Gene list used in this study.

**Table S2** 46 core landbird taxa under gene capture in this study.

**Table S3** Gene sequences cited from GenBank.

**Table S4** Positively selected genes identified by branch model based on the core landbird dataset.

**Table S5** Positively selected genes identified by branch-site model based on the core landbird dataset.

**Table S6** Function description of digestive system-related genes under adaptive evolution. Function description of genes is referred to NCBI Gene and/or OMIM and Kyoto Encyclopedia of Genes and Genomes (KEGG).

**Table S7** The genes under relatively relaxed selection (k < 1) and relatively intensified selection (k > 1) of our focal taxa (test branch) relative to the common ancestors of our focal taxa and their sister taxa (reference branch). This result is based on the core landbird dataset. Yellow, blue, and green show the genes involved in fat, protein, and carbohydrate digestion and absorption, respectively.

**Table S8** Digestive system gene sequences of Avian taxa cited from GenBank.

**Table S9** The genes under relatively relaxed selection (k < 1) and relatively intensified selection (k > 1) along the common ancestor branch (test branch) of core landbirds relative to the common ancestor branch (reference branch) of core landbirds and their sister taxa, waterbirds. This result is based on the whole avian dataset. Yellow, blue, and green show the genes involved in fat, protein, and carbohydrate digestion and absorption, respectively.

**Table S10** Positively selected genes identified by branch model based on the whole avian dataset.

**Table S11** Positively selected genes identified by branch site model based on the whole avian dataset. Yellow, blue, and green show the genes involved in fat, protein, and carbohydrate digestion and absorption, respectively.

