## Supplemental text for "Genomic bases underlying the adaptive radiation of core landbirds"

**Phototransduction pathway genes**

Lamb, T. D. Evolution of phototransduction, vertebrate photoreceptors and retina. Prog Retin Eye Res 36, 52-119 (2013).

Larhammar, D., Nordström, K. & Larsson, T. A. Evolution of vertebrate rod and cone phototransduction genes. Philos Trans R Soc Lond B Biol Sci 364, 2867-2880 (2009).

**Temperature-sensitive genes**

Dhaka, A., Viswanath, V. & Patapoutian, A. Trp ion channels and temperature sensation. Annu Rev Neurosci 29, 135-161 (2006).

Ferreira, G., Raddatz, N., Lorenzo, Y., González, C. & Latorre, R. in *TRP Channels in Sensory Transduction* (eds Madrid Rodolfo & Bacigalupo Juan) 1-39 (Springer, 2015).

Julius, D. TRP channels and pain. Annu Rev Cell Dev Biol 29, 355-384 (2013).

Latorre, R., Brauchi, S., Madrid, R. & Orio, P. A cool channel in cold transduction. Physiology 26, 273-285 (2011).

Liedtke, W. B. & Heller, S. *TRP Ion Channel Function in Sensory Transduction and Cellular Signaling Cascades*. (CRC Press, 2006).

McKemy, D. D. in *TRP Ion Channel Function in Sensory Transduction and Cellular Signaling Cascades*. (eds Wolfgang B Liedtke & Stefan Heller) 177-188 (CRC Press, 2006).

McKemy, D. D. The molecular and cellular basis of cold sensation. ACS Chem Neurosci 4, 238-247 (2013).

Patapoutian, A., Peier, A. M., Story, G. M. & Viswanath, V. ThermoTRP channels and beyond: mechanisms of temperature sensation. Nat Rev Neurosci 4, 529-539 (2003).

**Language-related genes**

Deriziotis, P. & Fisher, S. E. Neurogenomics of speech and language disorders: the road ahead. Genome Biol 14, 204, doi:10.1186/gb-2013-14-4-204 (2013).

Kang, C. & Drayna, D. Genetics of speech and language disorders. Annu Rev Genomics Hum Genet 12, 145-164 (2011).

Newbury, D. F. & Monaco, A. P. Genetic advances in the study of speech and language disorders. Neuron 68, 309-320 (2010).

Shimoyama, M. et al. The Rat Genome Database 2015: genomic, phenotypic and environmental variations and disease. Nucleic Acids Res 43, D743-D750 (2015).

**Hearing-related genes**

Crow, A. L. et al. The genetic architecture of hearing impairment in mice: evidence for frequency-specific genetic determinants. G3 (Bethesda) 5, 2329-2339 (2015).

Dandapat, A. et al. High frequency hearing loss and hyperactivity in DUX4 transgenic mice. PloS One 11, e0151467, doi:10.1371/journal.pone.0151467 (2016).

Fettiplace, R. & Hackney, C. M. The sensory and motor roles of auditory hair cells. Nat Rev Neurosci 7, 19-29 (2006).

Frucht, C. S., Uduman, M., Kleinstein, S. H., Santos-Sacchi, J. & Navaratnam, D. S. Gene expression gradients along the tonotopic axis of the chicken auditory epithelium. J Assoc Res Otolaryngol 12, 423-435 (2011).

Fuchs, P. A. How many proteins does it take to gate hair cell mechanotransduction? Proc Natl Acad Sci U S A 112, 1254-1255 (2015).

Gillespie, P. G. & Walker, R. G. Molecular basis of mechanosensory transduction. Nature 413, 194-202 (2001).

Jones, G., Teeling, E. & Rossiter, S. From the ultrasonic to the infrared: molecular evolution and the sensory biology of bats. Front Physiol 4, 117, doi:10.3389/fphys.2013.00117 (2013).

Keller, J. M. & Noben-Trauth, K. Genome-wide linkage analyses identify Hfhl1 and Hfhl3 with frequency-specific effects on the hearing spectrum of NIH Swiss mice. BMC Genet 13, 32 (2012).

Lavinsky, J. et al. Genome-wide association study identifies nox3 as a critical gene for susceptibility to noise-induced hearing loss. PLoS Genet 11, e1005094, doi:10.1371/journal.pgen.1005094 (2015).

Liu, Y. et al. The voltage-gated potassium channel subfamily KQT member 4 (KCNQ4) displays parallel evolution in echolocating bats. Mol Biol Evol 29, 1441-1450 (2012).

Liu, Z. et al. Parallel evolution of KCNQ4 in echolocating bats. PloS One 6, e26618, doi:10.1371/journal.pone.0026618 (2011).

Peng, A. W., Salles, F. T., Pan, B. & Ricci, A. J. Integrating the biophysical and molecular mechanisms of auditory hair cell mechanotransduction. Nat Commun 2, 523, doi:10.1038/ncomms1533 (2011).

Ru, B. et al. Molecular cloning and evolutionary analysis of GJB6 in mammals. Biochem Genet 50, 213-226 (2012).

Shen, B., Han, X., Jones, G., Rossiter, S. J. & Zhang, S. Adaptive evolution of the Myo6 gene in Old World fruit bats (Family: Pteropodidae). PloS One 8, e62307, doi:10.1371/journal.pone.0062307 (2013).

Smith, R., Shearer, A., Hildebrand, M. & Camp, G. in GeneReviews™ (eds Roberta A Pagon et al.) (University of Washington, 2008).

Steel, K. P. & Kros, C. J. A genetic approach to understanding auditory function. Nat Genet 27, 143-149 (2001).

Thiede, B. R. et al. Retinoic acid signalling regulates the development of tonotopically patterned hair cells in the chicken cochlea. Nat Commun 5, 3840, doi:10.1038/ncomms4840 (2014).

Zhao, B. et al. TMIE is an essential component of the mechanotransduction machinery of cochlear hair cells. Neuron 84, 954-967 (2014).

**Beak shape-related genes**

Abzhanov, A., Protas, M., Grant, B. R., Grant, P. R. & Tabin, C. J. Bmp4 and morphological variation of beaks in Darwin's finches. Science 305, 1462-1465 (2004).

Abzhanov, A. & Tabin, C. J. Shh and Fgf8 act synergistically to drive cartilage outgrowth during cranial development. Dev Biol 273, 134-148 (2004).

Bonneaud, C., Burnside, J. & Edwards, S. V. High-speed developments in avian genomics. *Bioscience* **58**, 587-595 (2008).

Brugmann, S. et al. Comparative gene expression analysis of avian embryonic facial structures reveals new candidates for human craniofacial disorders. Hum Mol Genet 19, 920-930 (2010).

Cheng, Y. et al. Evolution of beak morphology in the Ground Tit revealed by comparative transcriptomics. Front Zool 14, 58, doi:10.1186/s12983-017-0245-6 (2017).

Lamichhaney, S. et al. Evolution of Darwin’s finches and their beaks revealed by genome sequencing. Nature 518, 371-375 (2015).

Lamichhaney, S. et al. A beak size locus in Darwin’s finches facilitated character displacement during a drought. Science 352, 470-474 (2016).

**Protein, fat and carbohydrate digestion and absorption-related genes**

Kanehisa, M., Sato, Y., Kawashima, M., Furumichi, M. & Tanabe, M. KEGG as a reference resource for gene and protein annotation. Nucleic Acids Res 44, D457-D462 (2016).

Protein digestion and absorption pathway (KEGG-04974)

https://www.genome.jp/kegg-bin/show_pathway?map=hsa04974&show_description=show

Fat digestion and absorption pathway (KEGG-04975)

https://www.genome.jp/kegg-bin/show_pathway?map=hsa04975&show_description=show

Carbohydrate digestion and absorption pathway (KEGG-04973)

https://www.genome.jp/kegg-bin/show_pathway?map=hsa04973&show_description=show

**Taste transduction genes**

Kanehisa, M., Sato, Y., Kawashima, M., Furumichi, M. & Tanabe, M. KEGG as a reference resource for gene and protein annotation. Nucleic Acids Res 44, D457-D462 (2016).

Taste transduction pathway (KEGG-04742)

https://www.genome.jp/kegg-bin/show_pathway?map=hsa04742&show_description=show
