## Supplementary figures and images for "Genomic bases underlying the adaptive radiation of core landbirds"

### Supplemental Fig.1

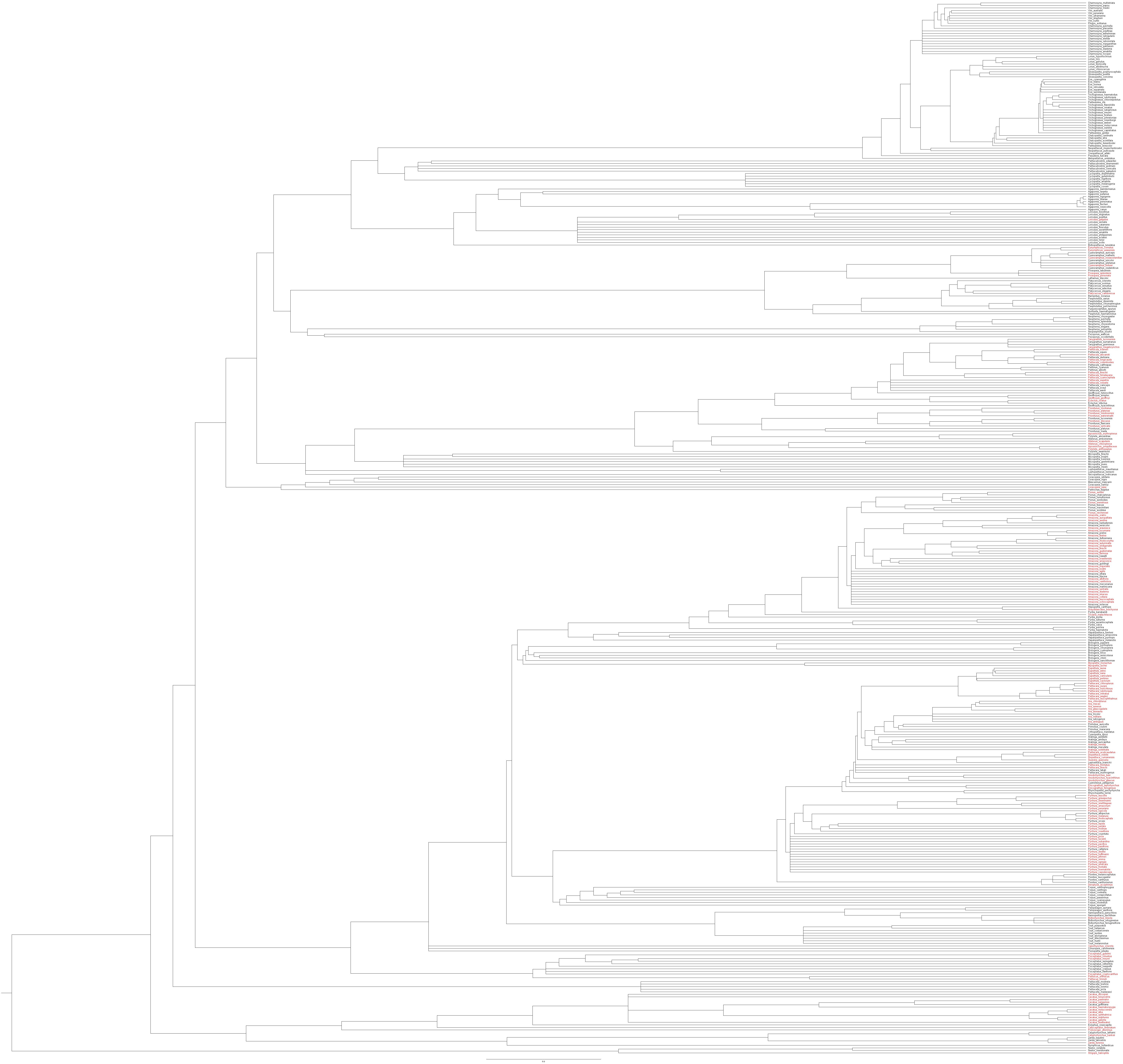
